# A Glycaemia-Prolonging Carbohydrate Gel Reduces Hunger and Cognitive Fatigue During Prolonged Esports Play

**DOI:** 10.64898/2026.01.06.697873

**Authors:** Takashi Matsui, Shion Takahashi, Takafumi Yamaguchi, Daisuke Funabashi, Takamasa Imada, Hiroki Shimizu

**Affiliations:** Institute of Health and Sport Sciences, University of Tsukuba, 1-1-1 Tennoudai, Tsukuba, Ibaraki 305-8574, Japan; R&D Center, Morinaga & Co., Ltd. 2-1-1 Shimosueyoshi, Tsurumi Ward, Yokohama City, Kanagawa 230-8504, Japan; Morinaga & Co., Ltd. 1-13-16 Shibaura, Minato City, Tokyo 105-8309, Japan

**Keywords:** esports, executive control, cognitive fatigue, hunger, continuous glucose monitoring, glycaemic stability, salivary cortisol

## Abstract

**Abstract:** Prolonged esports play can induce cognitive fatigue by placing sustained demands on executive control, decision-making, and stress regulation. Since the brain relies primarily on glucose as an immediate energy source, we hypothesized that a carbohydrate gel designed to prolong post-ingestion glycaemia stabilizes interstitial glucose, attenuates hunger and cognitive fatigue, and helps preserve performance during extended esports play. Eleven healthy young men completed a randomized, double-blind crossover study with two sessions in which they ingested one of two carbohydrate gels matched for total energy and total carbohydrate but differing in carbohydrate composition (high-fructose corn syrup [HFCS]-dominant vs maltodextrin [MDX]-dominant) before gameplay. Interstitial glucose was monitored continuously, while hunger, salivary cortisol, pupil diameter, executive function, and in-game performance were assessed repeatedly. Compared with the control (HFCS-dominant) condition, the glycaemia-prolonging formulation maintained higher interstitial glucose levels during the later phase of gameplay, blunted the progressive increase in hunger, and attenuated cortisol responses. These physiological and subjective advantages were accompanied by better executive control and more adaptive defensive behaviors in-game, including fewer fouls and more interceptions. In contrast, pupil diameter, an index of global arousal, remained unchanged across conditions, suggesting that benefits were not driven by altered arousal but by improved glycaemic stability and appetite regulation. These findings indicate that a carbohydrate formulation that prolongs glycaemia can reduce hunger-linked cognitive fatigue and support task-relevant behavior during prolonged esports play. This low-burden nutritional strategy may help sustain mental performance during prolonged digital activities that impose high cognitive demands.

**Highlights:**

- A glycaemia-prolonging gel maintained higher interstitial glucose late in esports play
- Hunger and salivary cortisol increased less with the glycaemia-prolonging formulation
- Executive control and defensive performance improved (fewer fouls, more interceptions)
- Pupil diameter was unchanged, suggesting benefits were not driven by altered arousal
- A low-burden, glycaemia-stabilizing strategy may help sustain cognitive stamina

## 1. Introduction

Esports, defined as organized competitive video gaming, has rapidly emerged as a major global sport with millions of active players worldwide and formal inclusion in international multi-sport events such as the Asian Games and the Olympic Esports Series (Jenny et al., 2017; Kemp et al., 2020; IOC, 2023; Qian et al., 2025). During prolonged matches, players must sustain high levels of attention, rapid decision-making, and fine motor control under time pressure, imposing substantial and sustained cognitive demands (Kemp et al., 2020; Matsui et al., 2024). In addition to its competitive aspect, prolonged esports play also models sustained digital knowledge work, in which individuals continuously process complex information, multitask, and interact with digital interfaces for extended periods (Kemp et al., 2020).

Recent laboratory work using a three-hour virtual football protocol has demonstrated that prolonged esports play induces declines in executive function accompanied by pupil constriction, indicating objective cognitive fatigue that can emerge even when subjective fatigue remains limited (Matsui et al., 2024). Using the same paradigm, changing the beverage consumed during play, replacing plain water with sparkling water, attenuated cognitive fatigue and preserved executive function over the three-hour session (Takahashi et al., 2025). These findings suggest that in-play ingestion can influence cognitive outcomes during prolonged esports.

Nutritional interventions that modulate postprandial glycaemia have been proposed as one approach to supporting cognition, because the brain depends primarily on glucose as a key energy substrate and cognitive performance can be sensitive to fluctuations in glucose availability (Álvarez-Bueno et al., 2019; Benton, 1990; Donohoe & Benton, 1999). Experimental studies have shown that acute increases in glucose availability can transiently enhance attention and other cognitive functions, particularly under cognitively demanding conditions. For example, ingestion of glucose tablet candy improves attentional performance after smartphone use (Setoguchi et al., 2025).

When the goal is sustained cognitive stamina rather than short-term facilitation, the temporal profile of post-ingestion glycaemia becomes critical. Carbohydrate type and processing shape the postprandial glycaemic profile. A more stable, sustained trajectory, often discussed in relation to glycaemic index (GI), has been associated with benefits for memory and attention over subsequent hours (Benton et al., 2003; Gaylor et al., 2022; Ingwersen et al., 2007; Micha et al., 2011; Philippou & Constantinou, 2014). These findings raise the possibility that promoting glycaemic stability may provide a sustained metabolic basis for prolonged cognitive performance, beyond simply increasing glucose availability acutely.

In addition to stabilizing glycaemia, carbohydrate formulations may influence cognitive stamina through effects on hunger and satiety. Dietary interventions that promote glycaemic stability are often accompanied by lower hunger (Emilien et al., 2017; Hafiz et al., 2022; Ischayek & Kern, 2006; White et al., 2020; Zafar et al., 2019; Zhu et al., 2021). Because hunger can bias attention and executive control toward food-related concerns and thereby compete with task-related goals (Loeber et al., 2013; Schiff et al., 2021), these appetite-related effects suggest a pathway by which glycaemic stability could support prolonged task performance by reducing hunger-related competition for cognitive resources during sustained mental effort.

However, most prior work has examined short-term effects on brief tasks, rather than tracking hunger, cognitive fatigue, and performance together across several hours. It therefore remains unclear whether shaping the postprandial glycaemic profile can produce a combined profile of attenuated hunger, reduced cognitive fatigue, and preserved performance during prolonged esports play—an ecologically relevant context in which players commonly postpone meals, experience progressively increasing hunger, and accumulate cognitive fatigue indexed by pupil constriction (Matsui et al., 2024). This pattern also suggests that carbohydrate-containing beverages or gels may be a practical vehicle for nutritional strategies targeting cognitive stamina in esports.

We therefore hypothesized that, compared with an energy-matched control formulation, a carbohydrate gel designed to prolong post-ingestion glycaemia would better stabilize interstitial glucose, attenuate hunger and cognitive fatigue, and help preserve performance during extended esports play. To test this hypothesis, we conducted a randomized, double-blind, crossover trial in which healthy young male casual esports players completed two three-hour virtual football sessions while consuming either a glycaemia-prolonging or a control carbohydrate gel matched for energy content. We monitored interstitial glucose continuously and repeatedly assessed hunger, executive function, in-game performance, and salivary cortisol.

## 2. Materials and Methods

### 2.1 Participants

Eleven healthy young adult men who casually enjoyed esports participated in this study. All participants were recruited from the local university community and surrounding area and were regular casual players of virtual football, but not professional esports athletes. Participant characteristics are shown in Table 1. The mean ± standard errors (SEM) age of the sample was 25.7 ± 1.1 years. Participants were non-smokers, reported no history of neurological, psychiatric, cardiovascular, metabolic, or endocrine disorders, and were not taking medications known to affect glucose metabolism or cognitive function. According to Matsui et al. (2024), casual players were defined as those who did not meet any of the following criteria: “participates regularly in virtual football competitions to win,” “belongs to an esports team, whether professional or amateur,” “has an in-game rank in the top 10th percentile or higher,” and “plays virtual football for more than 3 hours a day on average.”

**Table 1.** Participant characteristics of playing lifestyles.

|  |  | n = 11 |  |
| --- | --- | --- | --- |
| Age | Years | 25.7 | ± 1.1 |
| Sex | Male/female | 11 | / 0 |
| Weight | kg | 71.8 | ± 3.1 |
| Height | m | 1.8 | ± 0.0 |
| BMI | kg/m<sup>2</sup> | 23.1 | ± 0.7 |
| Playing career of proficient title | months | 6.6 | ± 1.5 |
| In-game rank | top percentile | 73.6 | ± 7.6 |
| Weekday esports playing time | min/day | 72.0 | ± 24.6 |
| Weekend esports playing time | min/day | 138.0 | ± 38.0 |
| Sleeping time | min/day | 408.0 | ± 12.0 |
| Vigorous-intensity activity time | min/day | 73.5 | ± 24.2 |
| Moderate-intensity activity time | min/day | 55.0 | ± 22.0 |
| Walking time | min/day | 33.0 | ± 6.16 |
| Exercise time | min/day | 42.0 | ± 22.6 |
| Sitting time | min/day | 486.0 | ± 107.5 |

Data are shown as mean ± SEM. Sex data are presented as the number of male and female participants. The in-game rank data show percentages from the highest to the lowest rank, ranging from 0 to 100.

We conducted an a priori power analysis using G*Power 3.1.9.7, assuming an effect size of Cohen’s f = 0.27 based on a previous study that demonstrated cognitive fatigue in casual esports players (Matsui et al., 2024). The analysis indicated that a total of 11 participants would be sufficient to detect a significant condition × time interaction (within-subjects) in a repeated-measures ANOVA at an alpha level of .05 and a statistical power (1 – β) of .80, with two conditions, four repeated measurements, a correlation among repeated measures of .50, and a non-sphericity correction ε of 1 (Faul et al., 2007). Eligibility criteria included age between 18 and 35 years, capacity to understand and sign informed consent, normal or corrected-to-normal vision sufficient for video game play, normal color vision, and availability to attend the experiment in person. Ethical approval for this study was granted by the Research Ethics Board of the University of Tsukuba, and the procedures complied with the Declaration of Helsinki.

### 2.2 Test Beverages

Two carbohydrate-containing gel beverages were used as test products in this study. Both beverages were developed by Morinaga & Co., Ltd. (Tokyo, Japan) and were provided in flexible pouch packaging suitable for single consumption. The two formulations were matched for total energy content and macronutrient composition, with identical volumes and similar sensory characteristics, but differed in the composition of their carbohydrate fraction.

Specifically, the control gel contained carbohydrates derived from high-fructose corn syrup (HFCS), whereas in the alternative formulation this HFCS-derived carbohydrates was replaced by maltodextrin (MDX) (Table 2). These formulations were developed with the aim of producing different post-ingestion glycaemic profiles (i.e., a faster rise/decline vs. a more sustained glycaemic maintenance).

**Table 2.** Nutritional composition of the control (HFCS-dominant) and prolonged-glycaemia (MDX-dominant) gel beverages (per pouch).

| Component | Control gel | Prolonged-glycaemia gel |
| --- | --- | --- |
| Energy (kcal) | 210 | 210 |
| Protein (g) | 0.0 | 0.0 |
| Fat (g) | 0.0 | 0.0 |
| Carbohydrate (g) | 52.4 | 52.4 |
| - maltodextrin (g) | 21.0 | 50.0 |
| - derived from high-fructose corn syrup (g) | 29.0 | 0.0 |
| - other ingredients (e.g., citric acid) (g) | 2.0 | 2.0 |
| - dietary fibre (g) | 0.4 | 0.4 |
| Salt equivalent (g) | 0.24 | 0.24 |

In the present manuscript, for readability, we refer to these beverages as the control (HFCS-dominant) gel and the prolonged-glycaemia (MDX-dominant) gel as an operational naming convention. Per pouch, both gel beverages provided 210 kcal, 0 g protein, 0 g fat, and 52.4 g of total carbohydrate, including 0.4 g dietary fibre, as well as 0.24 g of salt equivalent (Table 2). Thus, the only intended difference between the two products was the internal composition of the carbohydrate fraction.

The gel beverages were prepared and coded by the manufacturer and were supplied to the investigators in indistinguishable packages (Figure 1A). Allocation codes were concealed from both participants and experimenters until all data collection was completed, thereby maintaining the double-blind nature of the trial.

**Figure 1.**
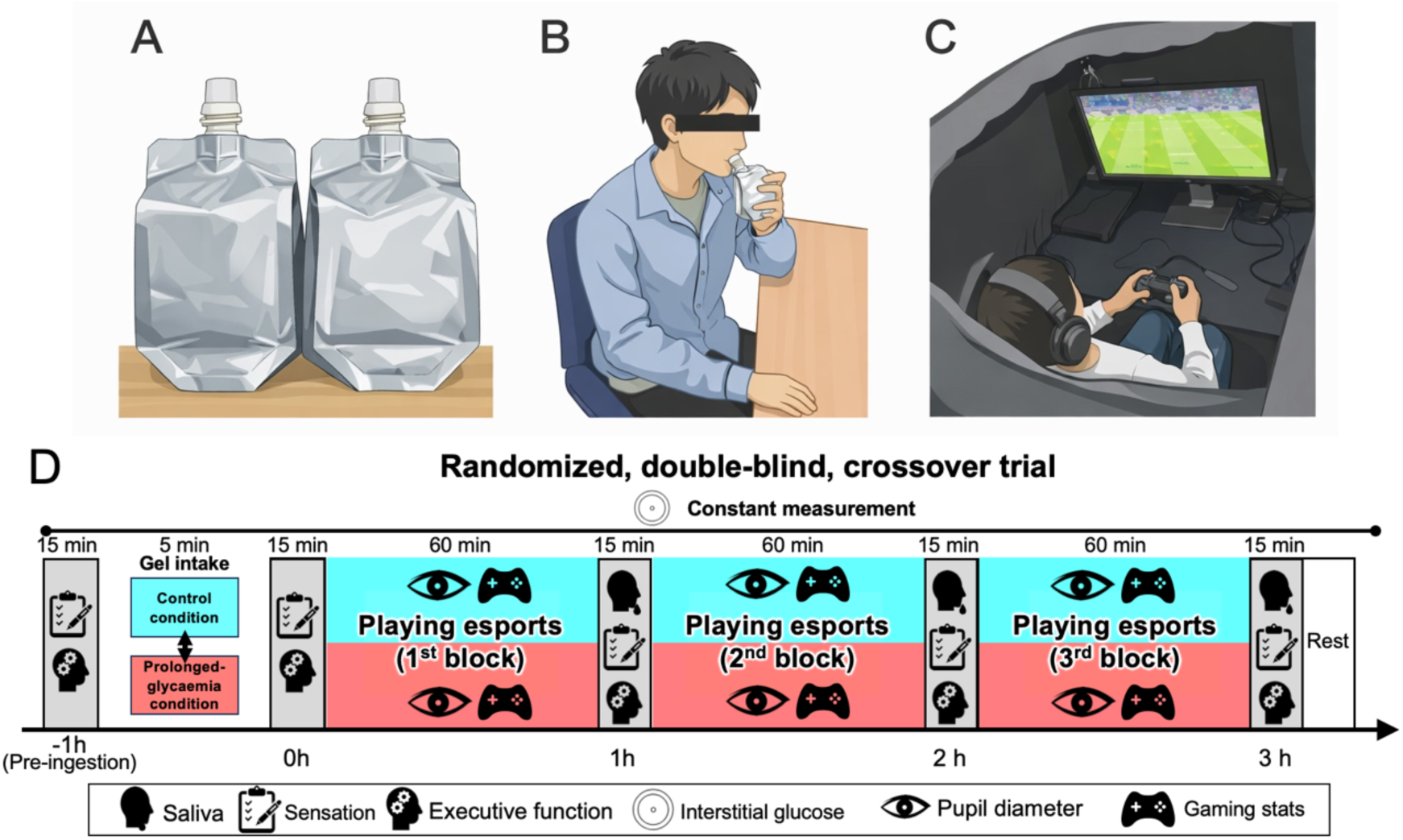
Experimental setup and timeline of the randomized, double-blind crossover protocol. (A) The two gel beverages (control [HFCS-dominant] vs prolonged-glycaemia [MDX-dominant]) were provided in identical pouches. (B) Representative image of gel intake. (C) Laboratory esports environment for virtual football play. (D) Timeline of the experiment. Participants completed two experimental sessions in a randomized, double-blind crossover design, separated by at least one week. On each visit, participants ingested the assigned gel 30 min before the start of gameplay; time 0 indicates gel ingestion. Participants then played virtual football for a total of 3 h, divided into three 60-min gameplay blocks separated by 15-min breaks for assessments. Subjective ratings, saliva sampling, and the executive function (flanker) task were administered during these breaks. Interstitial glucose was monitored continuously throughout the session using a continuous glucose monitoring system, and pupil diameter and in-game statistics were recorded during esports play. Time was referenced to gel ingestion, with −1 h indicating the pre-ingestion baseline assessment and 0 h indicating the assessment performed immediately after ingestion.

### 2.3 Study Design and Procedures

This study employed a randomized, double-blind, crossover design with two experimental sessions separated by at least one week. In each session, participants consumed one of two energy-matched carbohydrate gels: a control (HFCS-dominant) gel or a prolonged-glycaemia (MDX-dominant) gel (Figure 1). Condition order was counterbalanced using a computer-generated randomization sequence. The gels were supplied in indistinguishable packages and coded by the manufacturer; allocation codes were concealed from both participants and investigators until completion of data collection.

Two to seven days before the first experimental session, participants completed a familiarization visit (~60 min) under the same laboratory environment and game settings used in the main experiment to reduce learning effects. At the end of the familiarization visit, a continuous glucose monitoring (CGM) sensor (FreeStyle Libre Pro; Abbott, Japan) was fitted to the upper arm.

Before each experimental session, participants were instructed to complete their usual evening meal by 21:00 on the prior day, abstain from alcohol and vigorous physical activity from the prior day, and arrive in the morning after an overnight fast without breakfast or other energy intake. They were also asked to avoid caffeine-containing foods and beverages on the test day.

On each test day, participants first rested quietly upon arrival. Time was referenced to gel ingestion (0 h), with 0 h defined as immediately after ingestion. Baseline assessments were obtained 1 h before ingestion (−1 h), including subjective ratings (hunger and related sensations) and the flanker task. Participants subsequently ingested the assigned gel approximately 30 min before the start of the esports task (Figure 1B). Immediately after ingestion (0 h), subjective ratings were repeated and the flanker task was administered again.

Participants then performed three consecutive 60-min blocks of virtual football play separated by 15-min breaks for assessments (Figure 1C). During each break, saliva sampling, subjective ratings, and the flanker task were conducted at 1 h, 2 h, and 3 h after ingestion. Interstitial glucose was recorded continuously via CGM throughout the visit (Figure 1D). Participants were allowed to drink plain water ad libitum but were instructed to avoid any additional food or caloric beverages.

After completion of the final assessments, the session was terminated and participants were debriefed. The same procedures were repeated for the second session under the alternate gel condition. After the second session, the CGM sensor was removed.

### 2.4 Esports Play

The esports task consisted of playing a virtual football game from the eFootball series (Konami Digital Entertainment, Tokyo, Japan) continuously for 3 h. Participants played offline matches against a computer-controlled opponent throughout each session. Game settings were standardized and held constant across conditions and visits, including match mode, opponent difficulty, and camera view. Match duration was set to allow multiple full matches to be completed within each 1-h block.

Participants were seated at a fixed distance from the display and used a standard game controller. They were instructed to play as competitively and consistently as possible throughout the session, maintaining their usual tactics and formations and avoiding deliberate changes in play style between visits. Verbal encouragement was minimized to reduce external influences on motivation.

In-game performance metrics were extracted from match logs after each session. Outcomes included offensive and defensive indicators such as goals scored and conceded, total shots, shots on target, ball possession, pass success rate, fouls committed, and interceptions (pass cuts). For analysis, match-level values were averaged within each 1-h block (1 h, 2 h, 3 h) to yield participant-level hourly performance measures.

### 2.5 Measurements

#### 2.5.1 Interstitial Glucose Monitoring

Interstitial glucose concentration was monitored continuously using a professional continuous glucose monitoring system (FreeStyle Libre Pro; Abbott, Japan) attached to the upper arm. The sensor was inserted and initialized according to the manufacturer’s instructions during the familiarization visit (2–7 days before the first session), and data were recorded from pre-ingestion baseline through the end of the three-hour virtual football task.

Glucose values were exported at 15-min intervals. For time-series analyses, interstitial glucose was analysed across the pre-ingestion baseline and the 3-h gameplay period (from −5 h to +3 h relative to gel ingestion, where 0 h indicates immediately after ingestion) using a condition × time repeated-measures model. The pre-ingestion period from −5 h to 0 h represents the CGM-derived baseline before gel intake on the test day. Because CGM data were recorded continuously outside the laboratory, the −5 h to 0 h window includes ambulatory baseline data prior to arrival and pre-test procedures on the test day.

#### 2.5.2 Subjective Ratings (Hunger and Related Sensations)

Subjective sensations were assessed using 100-mm visual analogue scales (VAS), on which participants indicated their current state by placing a mark between “not at all” (0) and “extremely” (100). Ratings were obtained for hunger, fatigue, and enjoyment of play, as well as for additional sensations such as sleepiness, thirst, and urge to urinate as exploratory measures. VAS ratings were collected at five time points in each session: immediately before gel ingestion (pre-ingestion), immediately after ingestion, and 1 h, 2 h, and 3 h after ingestion during virtual football play. For longitudinal analyses, baseline-corrected (Δ) VAS values indicate change from the pre-ingestion baseline assessment at −1 h.

#### 2.5.3 Executive Function Task (Flanker)

Executive function was assessed during and after esports play using a computerized flanker task implemented in PsychoPy3 (v2021.1.2) (Peirce et al., 2019), with identical task parameters to those used in our previous study that successfully detected cognitive fatigue induced by prolonged virtual football play (Matsui et al., 2024). The task was executed on a desktop PC (Windows 10; Intel Core i7-11700 CPU; NVIDIA GeForce RTX 3070 Ti GPU; 16 GB × 2 RAM) and presented on a 23.8-inch LCD monitor (60 Hz refresh rate; 1920 × 1080 pixels). On each trial, a central target arrow was flanked by two arrows on each side, pointing either in the same direction (congruent trials) or the opposite direction (incongruent trials). Participants were instructed to respond as quickly and accurately as possible to the direction of the central arrow by pressing the corresponding key on a response device. For each test, stimuli were presented in a fully randomized order with 32 congruent and 32 incongruent trials (64 trials in total). Each trial consisted of a fixation cross (250 ms), stimulus presentation (up to 2000 ms), and a blank screen (1000 ms). The stimulus disappeared upon keypress and was followed by the blank screen until the next trial.

To reduce learning effects on cognitive performance, participants completed three practice sets of the task (192 trials in total), distributed across two occasions: on the day of the preliminary virtual football training session and immediately before the pre-test administration of the flanker task (Rappaport et al., 2025). Reaction times (RTs) and accuracy were recorded for all trials. Trials with RTs shorter than 200 ms or longer than 1500 ms were excluded as anticipations or lapses. Mean RT and percentage of correct responses were calculated separately for congruent and incongruent trials. The interference effect was quantified as the RT difference between incongruent and congruent trials. In addition, an inverse efficiency score (IES) was computed for the interference by dividing mean RT by the proportion of correct responses, providing a composite index of the speed–accuracy trade-off. For longitudinal analyses, Δ values indicate change from the pre-ingestion baseline assessment at −1 h. The flanker task was administered at −1 h (baseline, before gel ingestion), 0 h (immediately after gel ingestion), and at 1 h, 2 h, and 3 h after gel ingestion in each session.

#### 2.5.4 Esports Performance Outcomes

Esports performance was quantified using in-game statistics extracted from virtual football match logs. For each session, the following variables were recorded for every match: goals scored, goals conceded, total shots, shots on target, ball possession percentage, passes, number of interceptions (pass cuts), fouls committed. For analysis, match-level values were averaged within each 1-h block (1 h, 2 h, 3 h) to yield participant-level hourly performance measures (i.e., repeated measures across time). Particular attention was paid to fouls committed and interceptions during the later phase of play (2–3 h), as defensive indicators that are sensitive to cognitive fatigue during prolonged virtual football play.

#### 2.5.5 Salivary Cortisol

Salivary cortisol levels were measured as a physiological marker of stress associated with prolonged esports play. Saliva samples were collected at 1 h, 2 h, and 3 h after gel ingestion during the assessment breaks between gameplay blocks. At each time point, participants provided saliva by spitting directly into sterile collection tubes through a straw. Approximately 2 mL of saliva were collected at each sampling point and immediately stored at −80 °C. To precipitate mucus, tubes were later centrifuged at 1500 × g for 20 min, and the supernatant was aliquoted and stored at −80 °C until assay. Cortisol concentrations were determined using a commercially available enzyme-linked immunosorbent assay kit (Salivary Cortisol ELISA kit; Salimetrics, LLC, State College, PA, USA) according to the manufacturer’s instructions. All samples were measured in duplicate, and absorbance was read with a microplate reader (Varioskan LUX Multimode Microplate Reader; Thermo Fisher Scientific, Waltham, MA, USA). Cortisol concentrations were calculated from standard curves, and duplicate values were averaged for statistical analyses.

#### 2.5.6 Pupil Diameter

Pupil diameter was recorded as a primary index of central arousal and cognitive load during esports play, following the protocol described by Matsui et al. (2024). Pupil size was measured using an infrared eye tracker (Tobii Pro Nano; Tobii Technology, Sweden) during the entire three-hour virtual football task. Ambient lighting was standardized to 250–300 lux at eye level, and monitor brightness settings were kept constant across all measurements. Under these conditions, the visual stimuli presented during the experiment were expected to be comparable across participants, beverage conditions, and time points. The eye tracker was mounted below the display and connected to a dedicated laptop positioned next to the 23.8-inch monitor (60 Hz, 1920 × 1080 pixels). Participants were seated approximately 60 cm from the screen.

Before the first experimental session, a four-point calibration was performed using fixation targets presented at the four corners of the game screen. Following calibration, tracking accuracy was verified in Tobii Pro Lab (Tobii Technology, Sweden) by visually confirming that the recorded gaze position matched the instructed fixation targets. After successful calibration, binocular pupil diameter data were recorded continuously at a sampling rate of 60 Hz during all three-hour esports sessions. Pupil data were preprocessed in Tobii Pro Lab. Due to functional limitations of the device, eye blinks could not be explicitly detected. Short data losses of up to 80 ms were interpolated using the software’s built-in gap fill-in function. Noise reduction was then performed using the built-in moving median filter with a window size of five samples to smooth the signal. The final pupil diameter time series used for analysis was calculated as the average of the left and right eyes. To focus on broader trends in pupil diameter rather than second-to-second fluctuations, data were exported as 1-min averages, and hourly averages were subsequently computed for each of the three 1-h blocks (1 h, 2 h, 3 h). Because pupil diameter could only be recorded during gameplay, baseline measurements and z-normalization relative to pre-task levels were not performed. Only data segments with a capture rate of ≥80% during gameplay were included in the analyses.

### 2.6 Statistical analysis

All statistical analyses were performed using GraphPad Prism version 10 (GraphPad Software, San Diego, CA, USA) and R (version 4.x; R Foundation for Statistical Computing, Vienna, Austria). For interstitial glucose, the 3-h area under the curve (AUC) during virtual football play was compared between conditions using paired t-tests. For repeated-measures outcomes assessed across time (e.g., subjective ratings, flanker task indices, cortisol, and pupil diameter), condition (two levels) × time repeated-measures ANOVA were conducted. When a significant main effect or condition × time interaction was observed, planned pairwise comparisons between conditions at each time point were performed with Holm–Bonferroni correction for multiple testing. For in-game performance outcomes, match-level values were summarized as participant-level hourly averages (1 h, 2 h, 3 h) and analysed using condition × time repeated-measures models that respect the within-participant crossover structure (i.e., without treating match-level observations as independent). Data are presented as means ± SEM, and statistical significance was set at p < 0.05 (two-tailed). Effect sizes are reported as Cohen’s dz for paired comparisons and partial eta squared (η^2^_p_) for repeated-measures ANOVA. In addition, associations between ΔHunger and flanker indices at 2 h (ΔIES and ΔAccuracy for incongruent trials) were evaluated using Pearson’s correlation across observations pooled across conditions.

## 3. Results

### 3.1 Participant characteristics

All eleven eligible participants completed both experimental sessions and were included in the analyses. Participants were healthy young adult men who casually enjoyed esports (age: 25.7 ± 1.1 years, mean ± SEM; Table 1). No protocol deviations were recorded, and all experimental procedures were implemented as planned in both gel conditions.

### 3.2 Interstitial glucose

Interstitial glucose increased after ingestion of both gel beverages and peaked around the start of the virtual football task, then gradually declined over the subsequent three hours of play (Figure 2A). The time course differed between conditions: in the control (HFCS-dominant) condition, glucose declined more steeply during esports play and approached baseline values toward the end of the session, whereas in the prolonged-glycaemia (MDX-dominant) condition, glucose remained relatively elevated throughout the three-hour period.

**Figure 2.**
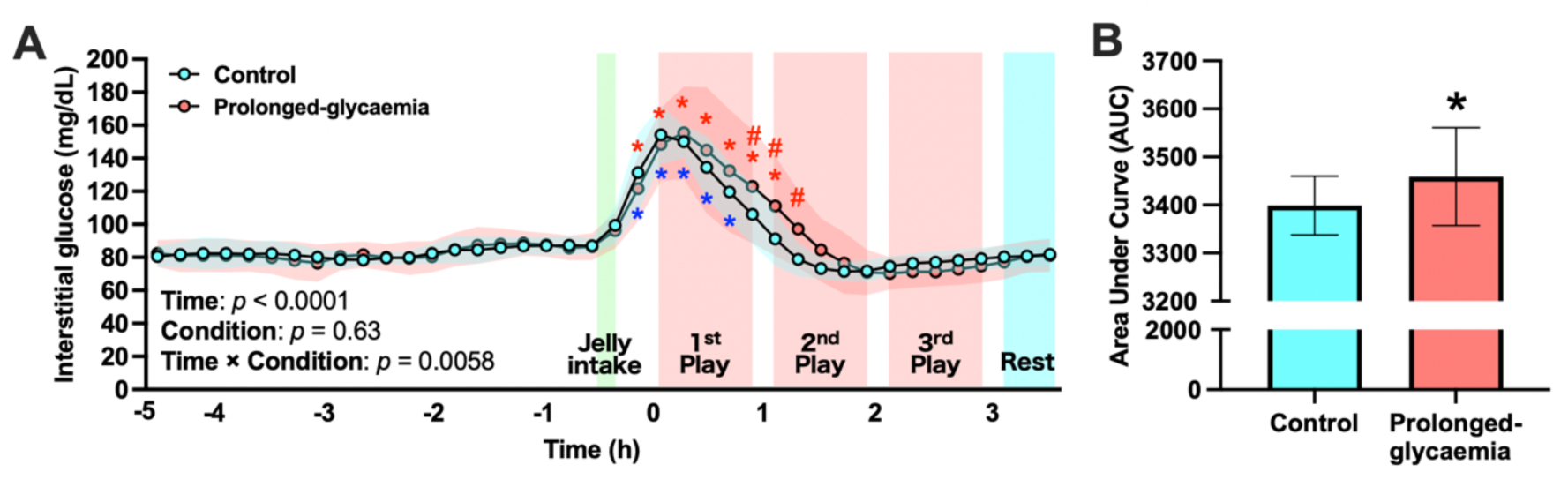
Interstitial glucose responses during prolonged esports play. (A) Interstitial glucose concentration (15-min samples) from −5 h to +3 h relative to gel ingestion; time 0 indicates immediately after gel ingestion. Values are shown as mean ± SEM. The shaded bands represent the SEM for the control (rapid-glycaemic; HFCS-dominant) and prolonged-glycaemia (MDX-dominant) conditions. Results of the condition × time repeated-measures analysis are shown in the lower-left corner of the graph (Time, Condition, and Interaction). Where indicated, post-hoc comparisons were performed with Holm–Bonferroni correction. *: *p* < 0.05 vs. −1 h (a stable pre-ingestion reference time point) within the same condition. #: *p* < 0.05 vs. control condition at the indicated time point. (B) Area under the curve (AUC) of interstitial glucose during the 3-h virtual football session. AUC was compared between conditions using paired t-tests. \**p* < 0.05 vs. control condition.

Using 15-min samples spanning the pre-ingestion baseline and the subsequent gameplay period (from −5.0 h to +3.5 h relative to gel ingestion; 0 h indicates immediately after ingestion), a condition × time repeated-measures ANOVA showed a significant main effect of time (F(39,312) = 43.48, p < 0.0001, η^2^_p_ = 0.845) and a significant time × condition interaction (F(39,295) = 1.741, p = 0.0058, η^2^_p_ = 0.187), whereas the main effect of condition was not significant (F(1,10) = 0.257, p = 0.626, η^2^_p_ = 0.031) (Figure 2A). Post-hoc comparisons were performed relative to a stable pre-ingestion reference time point (−1 h). For visualization, Figure 2A displays the time window from −5 h to +3 h.

Consistent with this pattern, the 3-h area under the curve (AUC) for interstitial glucose during virtual football play was greater in the prolonged-glycaemia condition than in the control condition (Figure 2B; p < 0.001, Cohen’s dz = 0.41), indicating more sustained glycaemic exposure over prolonged esports play.

### 3.3 Subjective ratings

Baseline (−1 h, pre-ingestion) ratings of hunger, sense of fatigue, and enjoyment did not differ between conditions. Immediately after gel ingestion, hunger decreased relative to pre-ingestion in both conditions, whereas fatigue and enjoyment showed no meaningful acute changes (Supplementary figure 1).

In the control condition, hunger increased progressively over the three hours of virtual football play, whereas this increase was attenuated in the prolonged-glycaemia condition (Figure 3A). A 2 (condition) × 4 (time: 0, 1, 2, 3 h) repeated-measures ANOVA on baseline-corrected hunger ratings showed a significant main effect of time [F(3,30) = 8.20, p = 0.00039, η^2^_p_ = 0.451] and a significant condition × time interaction [F(3,30) = 4.35, p = 0.0117, η^2^_p_ = 0.303], whereas the main effect of condition was not significant [F(1,10) = 1.48, p = 0.252, η^2^_p_ = 0.129]. Holm–Bonferroni–corrected post-hoc paired comparisons indicated that hunger was significantly lower in the prolonged-glycaemia than in the control condition at 2 h (p = 0.012, Cohen’s dz = 1.17), with no significant between-condition differences at 0, 1, or 3 h (p > 0.05; dz ≤ 0.24). Baseline-corrected hunger relative to −1 h (ΔHunger) is shown in Supplementary figure 2 for reference.

**Figure 3.**
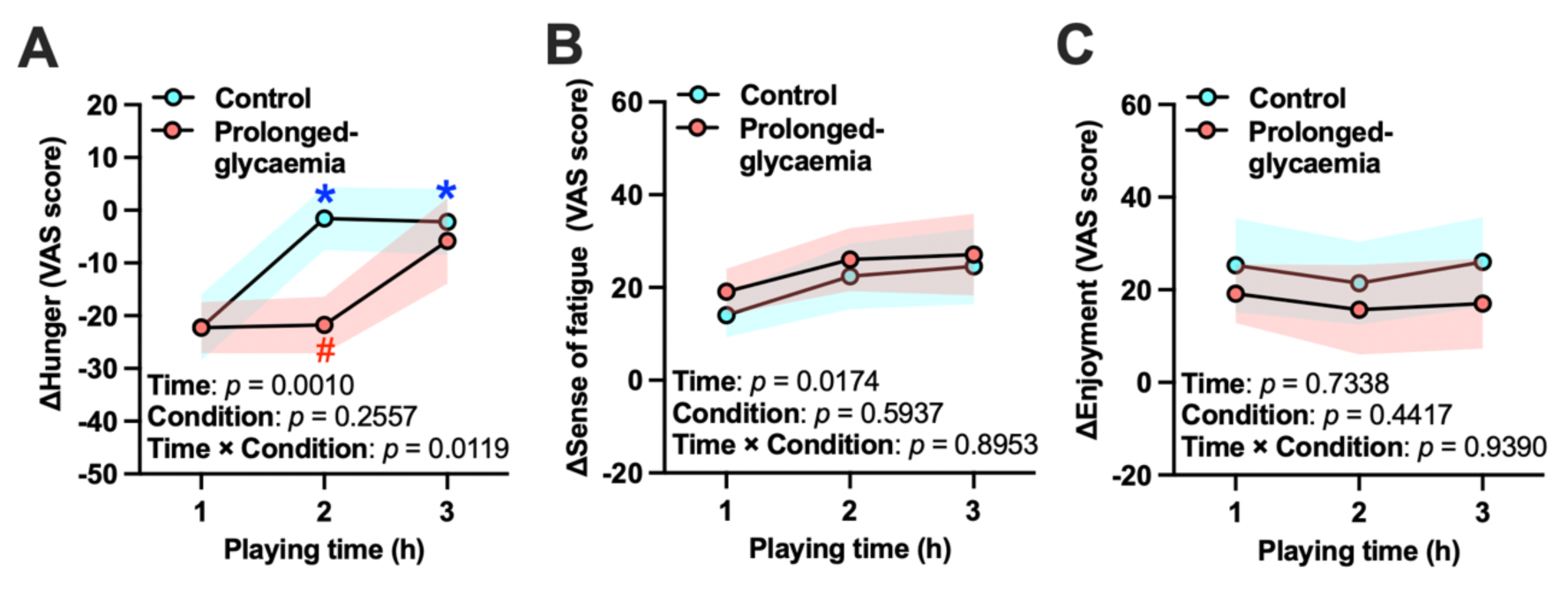
Subjective ratings during prolonged esports play. (A) Hunger ratings. (B) Fatigue ratings. (C) Enjoyment ratings. Values are shown as mean ± SEM. The shaded bands represent the SEM for the control (HFCS-dominant) and prolonged-glycaemia (MDX-dominant) conditions. Results of the condition × time repeated-measures ANOVA are shown in the lower-left corner of each graph (Time, Condition, and Interaction). Where indicated, post-hoc comparisons were performed with Holm–Bonferroni correction. \**p* < 0.05 vs. 0 h (immediately after gel intake) within the same condition. #*p* < 0.05 vs. control condition at the indicated time point. Baseline-corrected (Δ) values are expressed relative to the pre-ingestion baseline at −1 h (see Supplementary figure 2).

Ratings of fatigue and enjoyment of play exhibited main effects of time, with fatigue tending to increase and enjoyment remaining relatively high across the three-hour session, but no significant main effects of condition or condition × time interactions were observed (Figure 3B, C). For fatigue, baseline-corrected ratings showed a significant main effect of time (F(3,30) = 9.51, p = 0.0027, η^2^_p_ = 0.487), with no main effect of condition (F(1,10) = 0.8325, p = 0.446, η^2^_p_ = 0.059) and no condition × time interaction (F(3,30) = 0.29, p = 0.6619, η^2^_p_ = 0.028). Similarly, other subjective sensations such as sleepiness and urge to urinate showed no systematic differences between conditions. For enjoyment, there was a significant main effect of time (F(3,30) = 7.48, p = 0.0023, η^2^_p_ = 0.428), but no main effect of condition (F(1,10) = 0.73, p = 0.3259, η^2^_p_ = 0.068) and no condition × time interaction (F(3,30) = 0.07, p = 0.9381, η^2^_p_ = 0.007).

### 3.4 Executive function

Baseline flanker performance (−1 h, before gel ingestion) did not differ between conditions for any index. Flanker task indices did not change from pre-ingestion (−1 h) to immediately after ingestion (0 h) in either condition (Supplementary figure 3).

For all flanker indices, Δ values represent change from the pre-ingestion baseline at −1 h. Immediately after gel ingestion (0 h) and during prolonged virtual football play (1–3 h), ΔIES for incongruent trials tended to increase in the control condition, whereas they remained relatively stable in the prolonged-glycaemia condition (Figure 4A). A 2 (condition) × 4 (time: 0, 1, 2, 3 h) repeated-measures ANOVA on ΔIES showed a significant main effect of condition [F(1,10) = 5.47, p = 0.0414, η^2^_p_ = 0.354], with no significant main effect of time [F(3,30) = 0.29, p = 0.8496, η^2^_p_ = 0.028] and no condition × time interaction [F(3,30) = 1.00, p = 0.7101, η^2^_p_ = 0.091]. Overall, ΔIES was lower (better) in the prolonged-glycaemia than in the control condition across the session.

**Figure 4.**
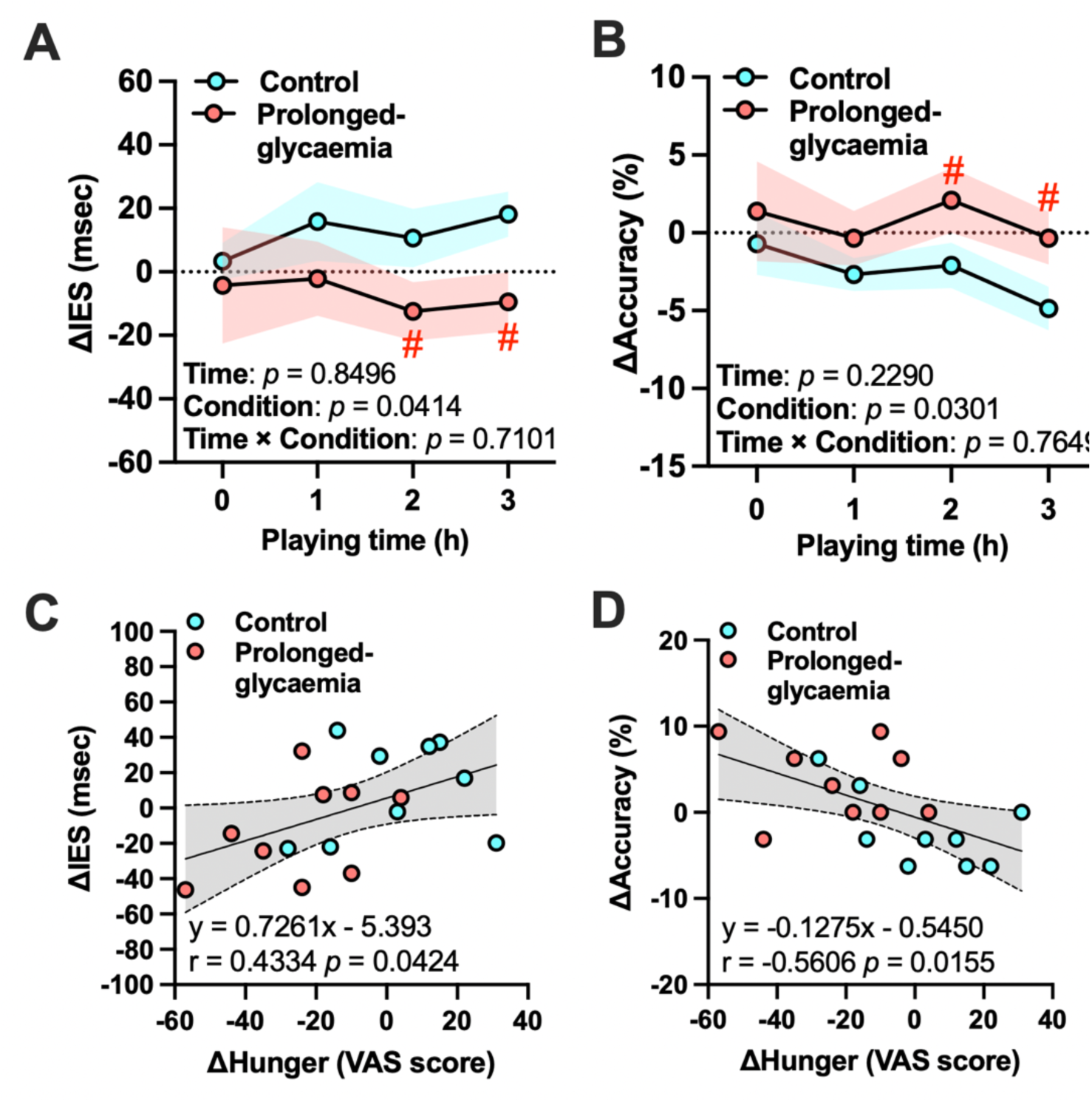
Executive function performance assessed by the flanker task during prolonged esports play. (A) ΔInverse efficiency score (IES) for interference. (B) ΔAccuracy (correct rate) for incongruent trials. Values are shown as mean ± SEM. Δ values indicate change from the pre-ingestion baseline assessment at −1 h. The shaded bands represent the SEM for the control (HFCS-dominant) and prolonged-glycaemia (MDX-dominant) conditions. Results of the condition × time repeated-measures ANOVA are shown in the lower-left corner of each graph (Time, Condition, and Interaction). Where indicated, post-hoc comparisons were performed with Holm–Bonferroni correction. \**p* < 0.05 vs. 0 h (immediately after gel intake) within the same condition. #*p* < 0.05 vs. control condition at the indicated time point. (C) Association between hunger ratings and ΔIES at 2 h. (D) Association between hunger ratings and Δaccuracy at 2 h. Scatter plots show individual observations at 2 h; colors indicate condition. The line and shaded band represent the fitted association and 95% confidence interval.

A similar pattern was observed for response accuracy on incongruent trials (Figure 4B). A repeated-measures ANOVA on Δaccuracy (Correct rate, %) showed a significant main effect of condition [F(1,10) = 6.60, p = 0.0301, η^2^_p_ = 0.398], with no significant main effect of time [F(3,30) = 0.69, p = 0.2290, η^2^_p_ = 0.064] and no condition × time interaction [F(3,30) = 0.97, p = 0.7649, η^2^_p_ = 0.088]. In contrast, mean reaction times and interference effects showed time-dependent changes without significant effects of condition. Taken together, these results indicate that the prolonged-glycaemia gel was associated with better executive control during prolonged esports play, particularly under the more demanding incongruent condition.

At 2 h of esports play, hunger ratings covaried with executive control (Figure 4C, D). Higher hunger was associated with higher ΔIES for incongruent trials (Figure 4C; r = 0.4334, p = 0.0424), and with lower Δaccuracy for incongruent trials (Figure 4D; r = −0.4664, p = 0.0410). These findings indicate that greater hunger at 2 h co-occurred with less efficient and less accurate executive performance.

### 3.5 Esports performance

Analyses of in-game virtual football performance revealed selective effects of the gel beverages on defensive but not offensive indicators (Figure 5). Offensive metrics, including goals scored, total shots, shots on target, and ball possession, did not differ significantly between conditions. For goals scored, a 2 (condition: control vs prolonged-glycaemia) × 3 (time: 1, 2, 3 h) repeated-measures ANOVA showed no main effect of condition (F(1,10) = 0.006, p = 0.9621, η^2^_p_ < 0.001) and no condition × time interaction (F(2,20) = 0.135, p = 0.7329, η^2^_p_ = 0.004), although a modest main effect of time was observed (F(2,20) = 4.45, p = 0.0031, η^2^_p_ = 0.107) (Figure 5A). Ball possession showed no effects of condition, time, or their interaction (condition: F(1,10) = 0.024, p = 0.7378, η^2^_p_ = 0.001; time: F(2,20) = 3.209, p = 0.1793, η^2^_p_ = 0.080; interaction: F(2,20) = 0.518, p = 0.7489, η^2^_p_ = 0.014) (Figure 5B). Shots on target showed the same pattern (condition: F(1,10) = 0.9012, p = 0.444, η^2^_p_ = 0.016; time: F(2,20) = 5.848, p = 0.0514, η^2^_p_ = 0.136; interaction: F(2,20) = 2.743, p = 0.0600, η^2^_p_ = 0.069) (Figure 5C). These findings indicate that the beverages did not meaningfully influence offensive production during prolonged esports performance.

**Figure 5.**
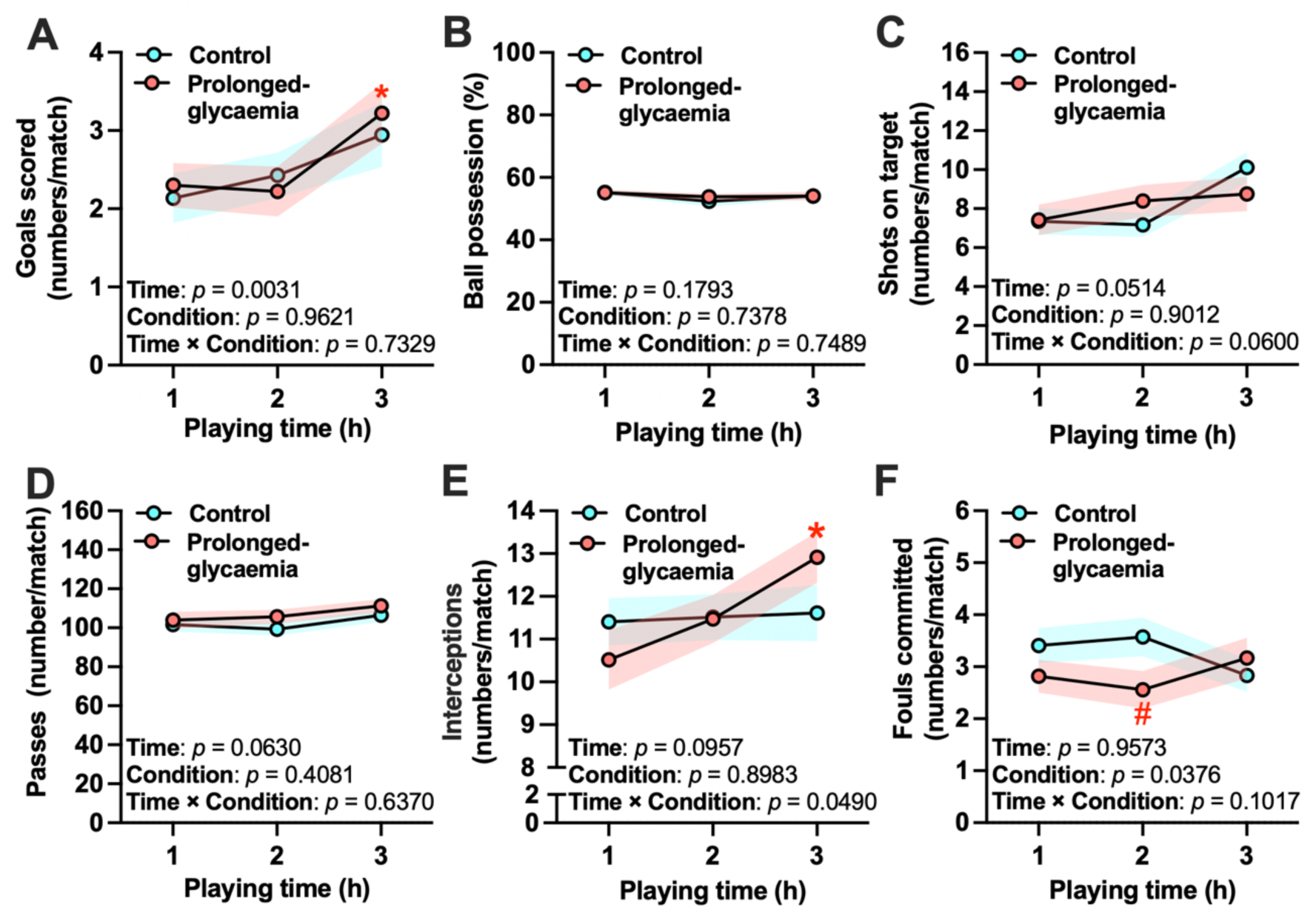
In-game virtual football performance during prolonged esports play. (A) Goals scored. (B) Ball possession. (C) Shots on target. (D) Passes. (E) Interceptions (pass cuts). (F) Fouls committed. Match-level values were extracted from game logs and summarized as participant-level hourly averages within each 1-h gameplay block (1 h, 2 h, 3 h). Values are shown as mean ± SEM. Results of the condition × time repeated-measures model are shown in the lower-left corner of each graph (Time, Condition, and Interaction). Where indicated, post-hoc comparisons between conditions at each time point were performed with Holm–Bonferroni correction. \**p* < 0.05 vs. 1st 1-h gameplay block (1 h) within the same condition. #*p* < 0.05 vs. control condition at the indicated time point.

In contrast, defensive performance metrics demonstrated clear condition-dependent patterns. Pass count did not show significant effects of condition, time, or their interaction (condition: F(1,10) = 0.382, p = 0.4081, η^2^_p_ = 0.010; time: F(2,20) = 1.721, p = 0.0630, η^2^_p_ = 0.044; condition × time: F(2,20) = 1.498, p = 0.6370, η^2^_p_ = 0.039) (Figure 5D). Interceptions (pass cuts) exhibited a significant condition × time interaction (F(2,20) = 4.434, p = 0.0490, η^2^_p_ = 0.107) (Figure 5E). Fouls committed also showed a significant main effect of condition (F(1,10) = 1.064, p = 0.0376, η^2^_p_ = 0.028) (Figure 5F). These results suggest that a carbohydrate formulation designed to prolong glycaemia selectively supports defensive decision-making and behavioral control during extended virtual football play.

### 3.6 Salivary cortisol

During prolonged virtual football play, cortisol levels increased across the three hours in the control condition, whereas this increase was attenuated in the prolonged-glycaemia condition (Figure 6A). A 2 (condition) × 3 (time: 1, 2, 3 h) repeated-measures ANOVA showed a significant main effect of condition [F(1,10) = 11.19, p = 0.0199, η^2^_p_ = 0.528] and a significant main effect of time [F(2,20) = 2.64, p = 0.0453, η^2^_p_ = 0.209], with no condition × time interaction [F(2,20) = 0.825, p = 0.2940, η^2^_p_ = 0.076]. Overall, cortisol concentrations were higher in the control than in the prolonged-glycaemia condition across the play period.

**Figure 6.**
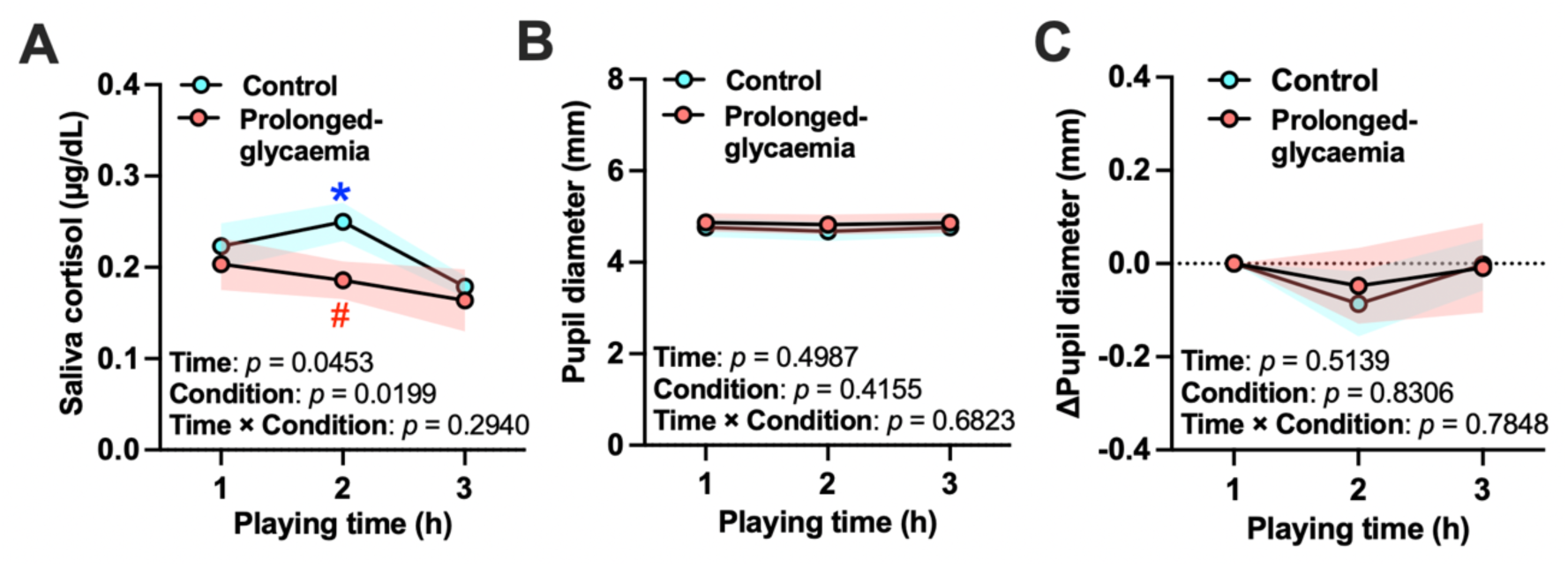
Physiological stress and arousal indices during prolonged esports play. (A) Salivary cortisol levels. (B) Pupil diameter. (C) Δ Pupil diameter. Values are shown as mean ± SEM. The shaded bands represent the SEM for the control (HFCS-dominant) and prolonged-glycaemia (MDX-dominant) conditions. Results of the condition × time repeated-measures ANOVA are shown in the lower-left corner of each graph (Time, Condition, and Interaction). Where indicated, post-hoc comparisons were performed with Holm–Bonferroni correction. \**p* < 0.05 vs. 1h within the same condition. #*p* < 0.05 vs. control condition at the indicated time point.

### 3.7 Pupil diameter

Pupil diameter remained stable across the three-hour virtual football task in both beverage conditions, showing no systematic decline over time and no differences between conditions (Figure 6B). A 2 (condition: control vs prolonged-glycaemia) × 3 (time: 1, 2, 3 h) repeated-measures ANOVA on hourly mean pupil diameter revealed no significant main effects of condition (F(1,10) = 0.491, p = 0.4155, η^2^_p_ = 0.052) or time (F(2,20) = 0.675, p = 0.4987, η^2^_p_ = 0.070), and no condition × time interaction (F(2,20) = 0.264, p = 0.6823, η^2^_p_ = 0.029). Similarly, no significant difference was observed in Δpupil diameter (Figure 6C). Thus, the gel conditions did not meaningfully influence pupil diameter during prolonged esports play.

These null findings contrast with our previous study, in which pupil diameter decreased progressively over three hours of esports play when participants had completed breakfast approximately two hours prior to testing (Matsui et al., 2024). In the present study, participants ingested a carbohydrate-containing gel beverage approximately 30 min before gameplay, which may have helped maintain central arousal or prefrontal activity during the early and mid phases of play, thereby preventing the gradual pupil constriction typically associated with cognitive fatigue.

## 4. Discussion

The present randomized, double-blind crossover study shows that promoting post-ingestion glycaemic stability can support cognitive endurance during prolonged esports play. Compared with an energy-matched control gel, the glycaemia-prolonging formulation maintained higher interstitial glucose during the later phase of gameplay (Figure 2), attenuated hunger (Figure 3), and was associated with better executive function (Figure 4) and more adaptive defensive behaviors (fewer fouls and more interceptions; Figure 5) with reduced cortisol responses (Figure 6). Pupil diameter did not differ between conditions, suggesting that these benefits were not driven by altered global arousal (Van Der Wel & Van Steenbergen, 2018). Together, these findings highlight postprandial glycaemic profile as a practical target for low-burden nutritional strategies in prolonged high-demand digital tasks.

### 4.1 Hunger as a Competing Cognitive Demand During Prolonged Esports Play

A central finding of the present study is that the prolonged-glycaemia gel attenuated the progressive rise in subjective hunger during prolonged esports play (Figure 2, 3), and that this pattern co-occurred with better executive function indices and more adaptive defensive in-game performance (Figure 4, 5). This observation aligns with evidence that hunger can act as a competing cognitive demand, biasing attentional and executive resources toward food-related processing and impairing inhibitory control even in tasks unrelated to food (Loeber et al., 2013; Montagrin et al., 2021; Schiff et al., 2021). Within this framework, rising hunger during extended cognitively demanding activity may draw upon limited cognitive resources, thereby reducing the capacity available for rapid response selection, strategic anticipation, and other executive processes required for esports performance.

Consistent with this interpretation, hunger increased more in the control condition and co-occurred with less favorable executive control indices, whereas the prolonged-glycaemia condition showed attenuated hunger with cognitive performance (Figure 4A, B). Supporting this link at the key 2-h time point, higher hunger was associated with worse incongruent-trial performance (higher ΔIES and lower accuracy; Figure 4C, D). Although correlational, these findings are consistent with the possibility that appetite-related signals compete for executive resources during sustained digital performance, contributing to hunger-linked cognitive fatigue during prolonged esports play.

### 4.2 Blood Glucose Stability as a Metabolic Basis for Sustained Cognitive Performance

Another factor that may underlie the observed benefits is the stabilization of interstitial glucose during prolonged esports play. Nutritional studies have shown that carbohydrate formulations shape the postprandial glycaemic profile, and that more stable or sustained glycaemic trajectories have been associated with benefits for memory, attention, and sustained mental effort several hours after ingestion (Benton et al., 2003; Gaylor et al., 2022; Philippou & Constantinou, 2014). The present findings are consistent with this framework: interstitial glucose remained relatively elevated during the later phase of the three-hour session in the prolonged-glycaemia condition (Figure 2), coinciding with better executive control indices and more adaptive defensive performance (Figure 4, 5).

Importantly, these effects were observed without detectable differences in pupil diameter (Figure 6B), suggesting that the gel formulation did not measurably alter global arousal. Instead, the results are consistent with the possibility that glycaemic stability supported task performance by providing a more stable metabolic milieu while simultaneously reducing subjective hunger that could otherwise compete for cognitive resources. This interpretation is in line with evidence that acute glucose intake can transiently benefit attention under cognitive load (Setoguchi et al., 2025), and further highlights the potential importance of the temporal glycaemic profile, rather than energy load alone, during prolonged, high-demand esports activity.

More broadly, the present findings support the view that cognitive endurance during complex digital tasks may depend on interactions between metabolic state, subjective energy-related signals, and executive control (Benton, 1990; Boksem & Tops, 2008; Meeusen, 2014). By promoting glycaemic stability and attenuating hunger, the prolonged-glycaemia gel may create conditions that favor sustained cognitive engagement without requiring an increase in baseline arousal.

### 4.3 Broader implications for modern cognitive work and real-world soccer performance

Beyond esports, these results may have implications for other forms of prolonged digital work that demand sustained attention, rapid decision-making, and continuous motor–cognitive coordination. The observation that a glycaemia-stabilizing nutritional strategy was associated with better executive control and task-relevant behavior without altering global arousal suggests that similar mechanisms could be relevant in occupational or daily settings that require prolonged cognitive engagement (Figure 4-6). In such contexts, stabilizing metabolic state and mitigating hunger-related cognitive interference may offer a practical approach to supporting cognitive stamina during extended periods of high workload.

The implications may also extend, cautiously, to real-world sports. The virtual football task captures key perceptual–cognitive demands relevant to association football, including continuous scanning, rapid pattern interpretation, prediction of opponents’ intentions, and time-pressured decisions. Prior research indicates that mental fatigue can impair these components in soccer players, affecting passing accuracy, defensive positioning, and decision-making speed after sustained cognitive load (Badin et al., 2016; Smith et al., 2016; Thompson et al., 2019). By showing that a gel formulation associated with more stable glycaemia and lower hunger was accompanied by better executive control and defensive behaviors in a football-like cognitive task (Figure 2-5), the present findings raise the possibility that nutritional approaches targeting glycaemic stability could complement existing strategies to manage cognitive fatigue in team sports. However, this extrapolation remains speculative because physical demands and real-world social coordination differ substantially between virtual and field-based environments.

### 4.4 Limitations and future directions

Several limitations should be acknowledged. First, the sample size was modest and consisted exclusively of healthy young adult men who were casual virtual football players, limiting generalizability to women, older adults, professional esports athletes, or individuals with metabolic or psychiatric conditions. Second, the task involved a single esports title performed in an offline, laboratory setting without concurrent physical load; it remains unclear whether similar benefits would be observed in other esports genres, during online competitive play, or under physically demanding conditions.

Third, the test products were developed by a single manufacturer, and only one dose and ingestion timing (approximately 30 min before play) were examined. Other formulations, energy contents, or dosing schedules (e.g., repeated intake during play) may yield different metabolic and cognitive patterns. In addition, the two gels differed in carbohydrate composition

(HFCS-dominant vs MDX-dominant), and the present study operationally interprets their effects through the observed glycaemic profile rather than a direct measurement of digestion/absorption kinetics. Future work should characterize formulation properties more directly (e.g., glycaemic index testing, in vitro digestion metrics, or gastric emptying-related measures) and examine whether other carbohydrate matrices that yield sustained glycaemic profiles produce similar effects.

Fourth, while interstitial glucose, hunger, cortisol, pupil diameter, executive function, and in-game behavior were measured concurrently, the analyses were primarily univariate and do not establish causal mediation pathways. Larger samples with multivariate or time-series approaches will be needed to test whether changes in hunger and/or cortisol statistically mediate the association between glycaemic stability and performance. Finally, pupil diameter could not be recorded at a true resting baseline, and only global arousal rather than regional brain activity was assessed. Combining pupillometry with neurophysiological methods (e.g., EEG or fNIRS) would provide a more detailed picture of the neural mechanisms through which nutritional interventions influence cognitive endurance.

Despite these limitations, the current study indicates that a carbohydrate gel associated with a more sustained glycaemic profile can attenuate hunger, reduce stress-related cortisol responses, and support executive control and defensive esports performance during prolonged play, without detectable changes in global arousal indexed by pupil diameter. These findings highlight the potential value of targeting post-ingestion glycaemic stability as a practical nutritional approach to support cognitive endurance in prolonged, high-demand esports and related modern cognitive activities.

## 5. Conclusion

Our findings support the hypothesis that, during extended esports activity, consuming a carbohydrate gel associated with a more sustained post-ingestion glycaemic profile can stabilize interstitial glucose, attenuate hunger-related cognitive fatigue, and support in-game performance. These effects occurred without changes in pupil diameter, suggesting that the benefits were not driven by altered global arousal but were consistent with improved glycaemic stability and appetite regulation during prolonged play. Because the two gels were matched for energy and total carbohydrate, the post-ingestion glycaemic profile may be more important than carbohydrate load per se for cognitive endurance. A low-burden, glycaemia-stabilizing nutritional strategy may therefore help sustain mental performance during prolonged digital activities that impose high cognitive demands.

## Data availability

The data supporting the findings of this study are available from the corresponding author upon request.

## Author contribution

T.M., T.I. and Y.S. conceived and designed the study. T.M., S.T., T.Y., and D.F. recruited participants, collected the data, conducted data analysis and interpreted data. T. M. drafted the manuscript. T.M., T.I. and Y.S. edited and revised the manuscript. All authors approved the final version.

---

**Supplementary figure 1.**
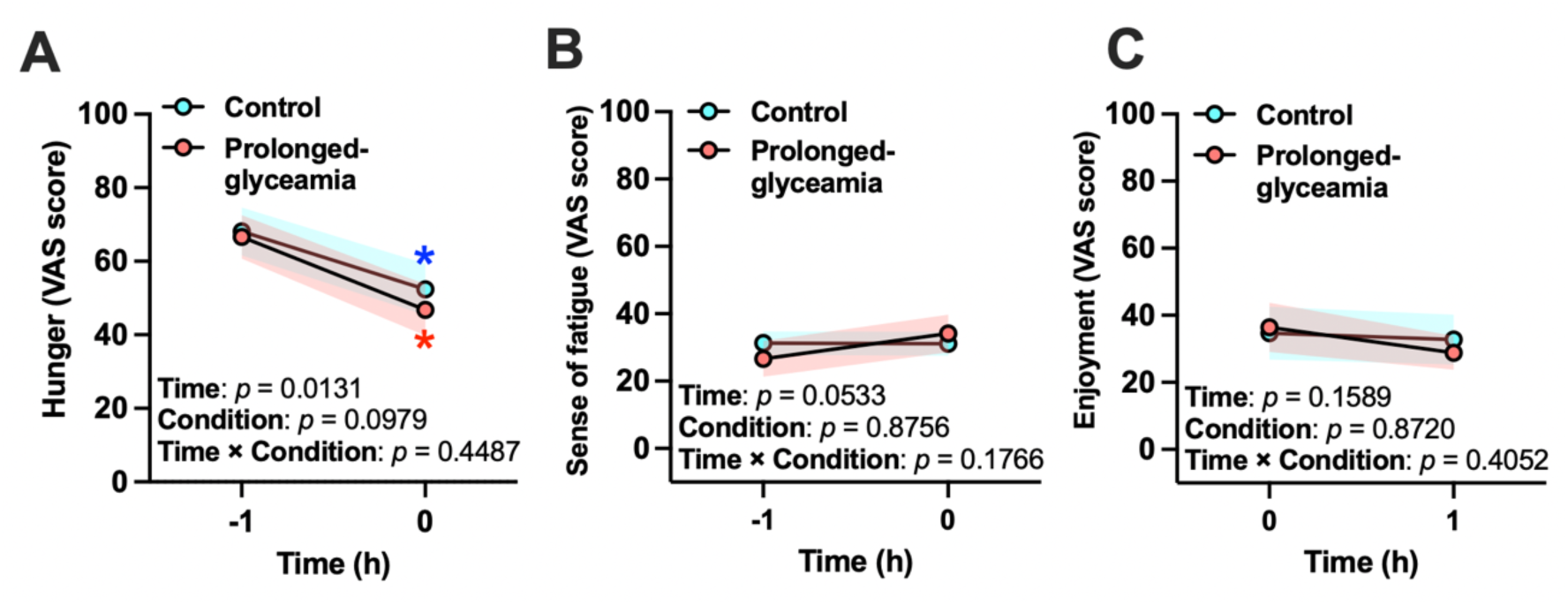
Subjective ratings before and immediately after gel ingestion. (A) Hunger. (B) Fatigue. (C) Enjoyment. Values are shown as mean ± SEM. Shaded bands indicate the SEM for the control (HFCS-dominant) and prolonged-glycaemia (MDX-dominant) conditions. Condition × time (pre vs immediately post ingestion) repeated-measures ANOVA results are shown within each panel (Time, Condition, and Interaction). Where indicated, Holm–Bonferroni–corrected post-hoc tests were used for within-condition comparisons between time points. *p < 0.05 vs. −1 h within the same condition.

**Supplementary figure 2.**
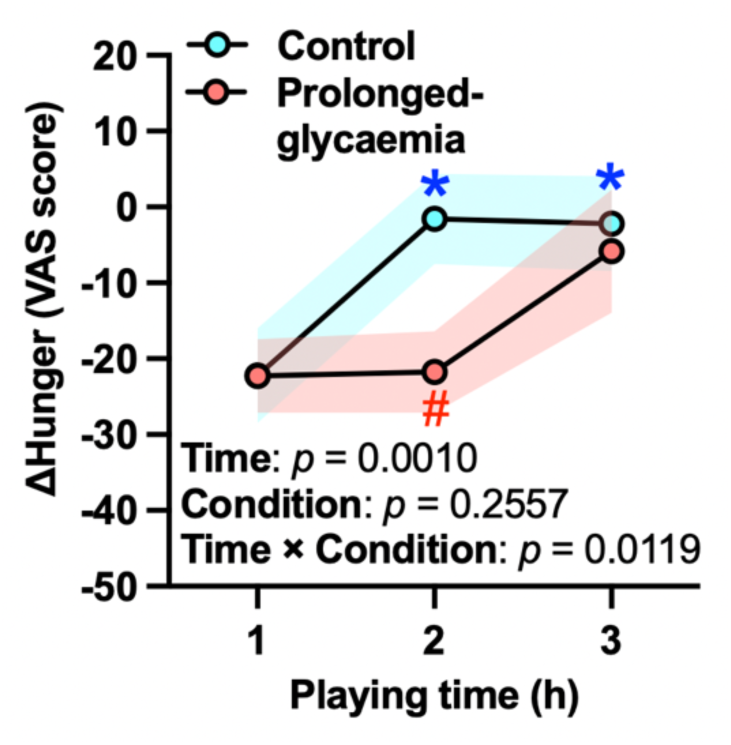
Baseline-corrected hunger during prolonged esports play. ΔHunger was calculated as the change in hunger VAS score relative to the pre-ingestion baseline at −1 h (Δ = value at each time point − value at −1 h). −1 h is used as the reference but is not plotted on the x-axis and negative values indicate lower hunger than baseline (−1 h). Data are shown at 1 h, 2 h, and 3 h after gel ingestion (0 h = immediately after ingestion) for the control (HFCS-dominant) and prolonged-glycaemia (MDX-dominant) conditions. Values are mean ± SEM. *p < 0.05 vs. 1 h within the same condition; #p < 0.05 vs. the control condition at the indicated time point (Holm–Bonferroni corrected).

**Supplementary figure 3.**
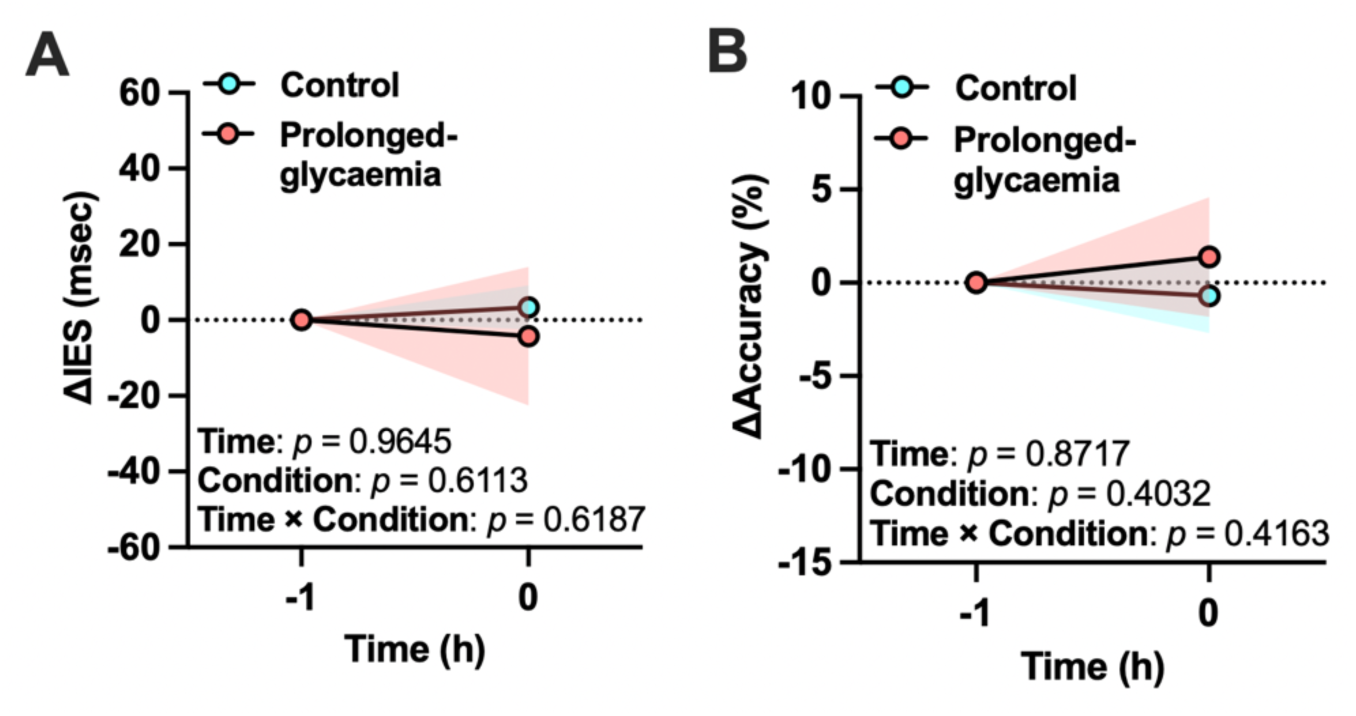
Executive function performance before and immediately after gel ingestion. (A) ΔInverse efficiency score (IES) for interference. (B) ΔAccuracy (correct rate) for incongruent trials. Values are shown as mean ± SEM. Shaded bands indicate the SEM for the control (HFCS-dominant) and prolonged-glycaemia (MDX-dominant) conditions. Condition × time (−1 h vs 0 h; pre vs immediately post ingestion) repeated-measures ANOVA results are shown within each panel (Time, Condition, and Interaction), indicating no detectable changes from −1 h to 0 h in either condition.

